# Overcoming Topology Bias and Cold-Start Limitations in Drug Repurposing: A Clinical-Outcome-Aligned LLM Framework

**DOI:** 10.64898/2026.01.12.699148

**Authors:** Ruihu Du, Mancheong Fung, Yuanjia Hu, Dongyang Liu

**Affiliations:** State Key Laboratory of Mechanism and Quality of Chinese Medicine, Institute of Chinese Medical Sciences, University of Macau, Macao SAR, China; Macafex LLC, 7 Stark Court, Ringoes, NJ 08551, USA; Centre for Pharmaceutical Regulatory Sciences, University of Macau, Macao SAR, China; DPM, Faculty of Health Sciences, University of Macau, Macao SAR, China; Drug Clinical Trial Center, Department of pharmacy, Peking University Third Hospital, Beijing, China; Center of Clinical Medical Research, Institute of Medical Innovation and Research, Peking University Third Hospital, Beijing, China; Beijing Key Laboratory of Cardiovascular Receptors Research, Peking University Third Hospital, Beijing, China

## Abstract

Graph Neural Networks (GNNs) in drug repurposing suffer from two limitations: transductive failure in zero-shot (cold-start) scenarios and popularity bias that misidentifies high-degree nodes as effective drugs. We propose a framework that shifts the optimization objective from graph topology to clinical utility, integrating Knowledge Graph RAG (KG-RAG), Supervised Fine-Tuning (SFT), and Kahneman-Tversky Optimization (KTO) using Phase III clinical trial outcomes as rewards. We evaluated our approach on a rigorous 1:10 negative sampling benchmark derived from MiRAGE, covering Standard, Cold-Start, and Degree-Matched settings. In Cold-Start scenarios where topological signals are absent, traditional GNNs (including TxGNN) collapse (Top-10 Precision < 0.30), whereas DR-SFT model achieves 0.80, demonstrating robust semantic generalization for novel compounds. Crucially, in Degree-Matched tests dominated by “popular but ineffective” decoys, the DR-KTO acts as a clinical gatekeeper, achieving 0.90 Top-10 Precision—significantly outperforming DR-SFT (0.70) and GNNs (0.2–0.4) by effectively penalizing hard negatives. Beyond repurposing accuracy, the model achieves state-of-the-art performance on BioASQ, and increased ability in Chemprot. Orthogonal validation via DrugReAlign confirms physical plausibility, yielding significantly lower docking binding energies for KTO-recommended candidates. SPR experiments further corroborate these findings. By aligning LLM reasoning with clinical evidence, our framework successfully bridges the gap between semantic inference, topological structure, and clinical reality.

## Introduction

Drug repurposing, the strategy of identifying new therapeutic indications for existing approved or investigational drugs, has emerged as a critical alternative to de novo drug discovery^1,2^. Compared to the traditional 10-15 years’ timeline and over $2 billion cost of developing a new molecular entity, re-purposing significantly reduces development risks and capital expenditure by leveraging established safety profiles and pharmacokinetic data^3^. In the face of public health emergencies or rare diseases with limited commercial incentives, re-purposing offers a pragmatic pathway to accelerate the translation of treatments from bench to bedside^4^. Historically, drug repurposing has relied on approaches such as pharmacological analysis, high-throughput screening, clinical retrospective studies, and bioinformatics—often guided more by serendipity^2^. However, the recent explosion of biomedical data has catalyzed a paradigm shift toward computational mining as a more deliberate and data-driven strategy^5^.

To systematize this process, Biomedical Knowledge Graphs (KGs) have become the standard substrate, encoding structured relations among drugs, diseases, genes, and pathways to enable link prediction^6^. Graph Neural Networks (GNNs) exploit these topologies to infer missing links with strong accuracy in transductive settings^7^. Formally, GNNs operate by aggregating information from local neighborhoods:

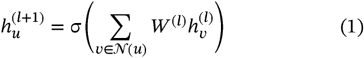

While effective for dense regions, this dependence on topology creates fundamental bottlenecks. First, as noted in recent scalability studies, GNNs often struggle with Zero-shot (Cold-Start) scenarios^8,9^. For a newly discovered disease or a novel compound *x*_*new*_ not present in the training graph, the neighbor set is empty (𝒩 (*x*_*new*_ ) = Ø ), causing the aggregation function to fail and predictive performance to collapse to random guessing^10^.

Second, GNNs suffer from Topology Bias (Popularity Bias). Current models typically maximize a likelihood function *P*(*edge*|*u,v*) that unintentionally correlates with node degree due to the power-law distribution of biological networks^11,12^ :

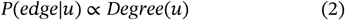

This results in “Hard Negatives”—well-studied drugs that appear to be perfect candidates due to their high connectivity but have failed in clinical reality. Although KGs curate diverse associations, they are often incomplete and slow to absorb new evidence, making purely structural inference brittle in dynamic therapeutic landscapes.

Parallel to structural approaches, Large Language Models (LLMs) have emerged as a transformative paradigm, shifting the focus from topological pattern matching to semantic reasoning. Unlike GNNs, which operate on the ontological structure of predefined graphs, LLMs leverage vast parametric knowledge to perform epistemological deduction, theoretically enabling generalization to unseen entities and mechanisms^13^. However, this semantic flexibility comes at a cost: without structural grounding, LLMs are prone to “hallucination,” generating plausible but factually incorrect associations due to the lack of explicit evidence^14,15^. To bridge the gap between structural fidelity and semantic reasoning, hybrid Retrieval-Augmented Generation (RAG) systems have been proposed. By injecting KG-derived paths into the prompt, these systems aim to ground Chain-of-Thought (CoT) reasoning in exp licit, multi-hop evidence^16–18^.

Yet, this integration introduces a subtle but critical vulnerability: the propagation of topological bias into the reasoning process. Since biomedical KGs follow a power-law distribution, retrieval mechanisms disproportionately surface “hub” nodes (popular, high-degree drugs) regardless of their specific causal relevance. Consequently, the generated CoT paths inevitably gravitate toward these popular entities, confounding popularity with causality. Crucially, when these topologically biased reasoning chains are subsequently used for Instruction Tuning (SFT)^19^, the bias is not merely retained but entrenched. The SFT process effectively teaches the model to memorize the graph’s structural imbalances, creating a “rich-get-richer” feedback loop where the model learns to mimic the stylistic form of reasoning while internalizing the heuristic that “high connectivity equals therapeutic efficacy,” rather than optimizing for genuine clinical truth^20^.

To dismantle this fallacy, we argue that structural likelihood (GNNs) and semantic plausibility (SFT) are necessary but insufficient; the ultimate arbiter must be clinical utility. While Preference Learning, typically Reinforcement Learning from Human Feedback (RLHF), has successfully aligned models in general domains^19,21^, it faces unique hurdles in biomedicine: it is prohibitively expensive to scale expert annotation, and crucially, even experts can be misled by plausible-sounding hallucinations (“sycophancy”)^22,23^. However, drug discovery offers a distinct advantage over open-ended chat: the existence of clinically anchored binary signals—Phase III trial successes vs. failures—which constitute objective, outcome-level supervision^24^. Leveraging this, we introduce a novel alignment mechanism using Kahneman–Tversky Optimization (KTO)^25^. Unlike methods requiring paired preference data, KTO is a lightweight framework that optimizes prospect-theoretic utility using unpaired binary rewards. By treating clinical outcomes as the ground truth “utility function,” KTO explicitly calculates a penalty gradient for “Hard Negatives”—candidates that are structurally central and semantically coherent but clinically futile. This effectively steers the model’s reasoning to distinguish between scientific hype (high degree) and therapeutic reality (proven efficacy).

Building upon this prospect-theoretic approach, we present a unified methodological framework that synergizes semantic reasoning with clinical value alignment. We formalize this pipeline into two progressive stages: DR-SFT (*Drug Repurposing via Supervised Fine-Tuning on the basis of Qwen3-8B*), which constructs a knowledge-graph-grounded reasoning engine to establish mechanistic plausibility; and DR-KTO (*Drug Repurposing via Kahneman-Tversky Optimization on the basis of DR-SFT* ), which refines this engine using historical clinical trial outcomes as the ultimate ground truth. Our primary objective is to transition computational drug discovery from a paradigm of *structural probability*—where candidates are selected based on data connectivity— to one of *clinical utility*, where decisions are guided by risk-aware alignment with human evidence. By bridging the epistemic gap between *in silico* predictions and *in vivo* realities, this framework not only mitigates the hallucinatory tendencies of generative models but establishes a robust, physically viable engine for identifying high-confidence therapeutic candidates in “cold-start” scenarios, ultimately accelerating the translation of computational insights into verified clinical assets.

## Results

### GNNs Struggle with Inductive Generalization and Hard Negatives

To systematically evaluate the limitations of topological approaches versus our semantic framework, we employed the MiRAGE^26^ benchmark, derived from the Comparative Toxicogenomics Database (CTD)^27^. We constructed a rigorous testing environment with a 1:10 positive-to-negative ratio to mimic the real-world scarcity of effective drugs. We compared our models against four graph baselines—RGCN, HGT, HAN, and TxGNN^9^ (re-implemented via Py torch Geometrics^28,29^)— across three distinct scenarios: Standard (relation-constrained), Cold-Start (inductive nodes), and Hard-Negative (degree-matched).

In the Standard setting (transductive link prediction), GNNs demonstrated strong performance, with TxGNN achieving competitive Precision-Recall metrics comparable to LLMs (Figure 2A). This confirms that GNNs are effective for completing missing links in well-structured graphs (Figure S2A–D). However, performance deteriorated sharply in Cold-Start scenarios (Figure 2C). Inductive limitations caused Top-10 Precision to plummet across all GNNs, with even the zero-shot-specialized TxGNN dropping to ∼0.30, while older GNNs approached zero (Figure S2H). Similarly, in the Hard-Negative setting designed to test resistance to popularity bias, GNNs failed to distinguish effective drugs from high-degree decoys, yielding consistently low Top-10 Precision (0.20–0.40) (Figure 2F; Figure S2L).

**Figure 1:**
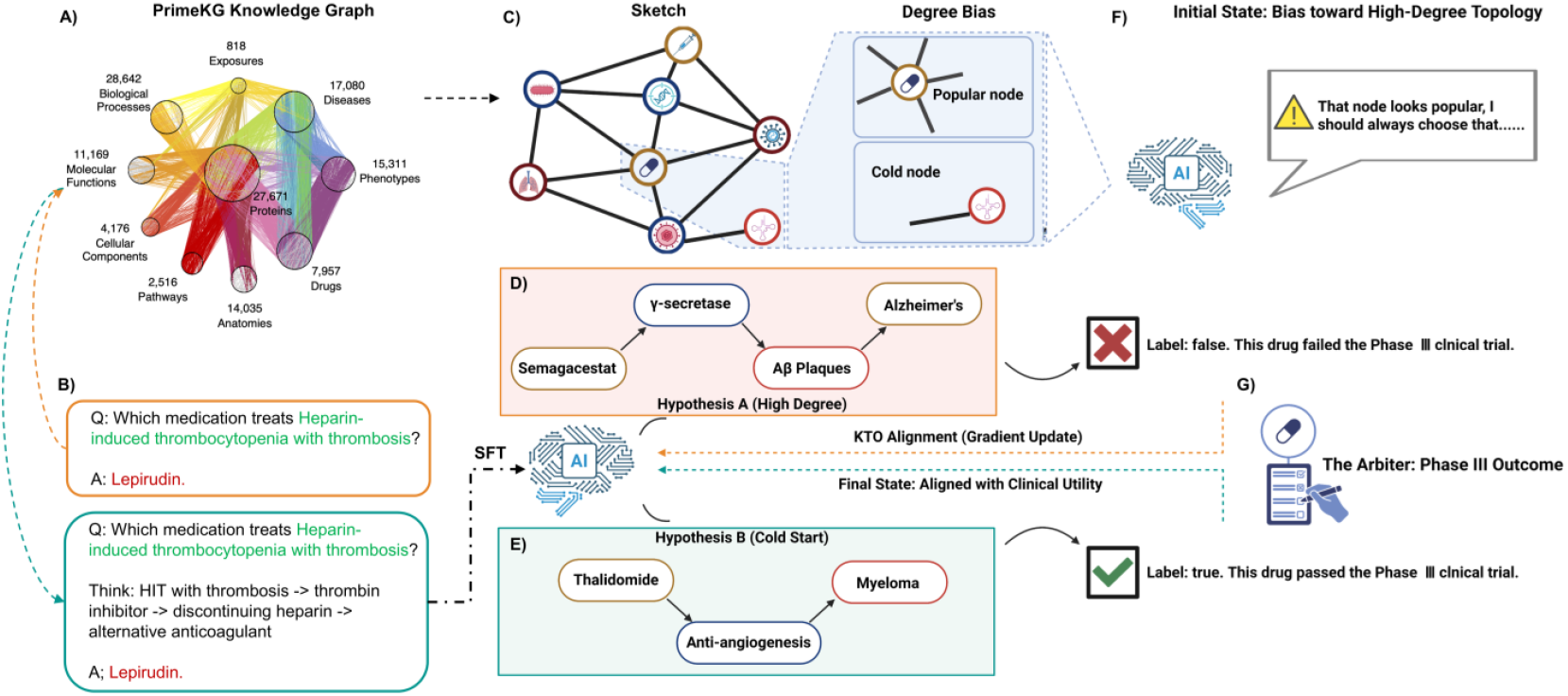
The Clinical-KTO Framework: Bridging Semantic Reasoning with Clinical Value Alignment. (A, B) Knowledge-Grounded Instruction Tuning. (A) The framework leverages PrimeKG, a comprehensive biomedical knowledge graph containing 17,080 diseases, 7,957 drugs and so on. (B) Raw QA pairs from RepoDB are enriched via KG-guided path retrieval to generate high-quality Chain-of-Thought (CoT) reasoning trajectories, which are used to fine-tune the base LLM (SFT stage). (C, F) The Topology Bias Challenge. (C) A schematic representation of the graph structure highlights the disparity between “hot” hub nodes (high-degree) and “cold” peripheral nodes. (F) Traditional models exhibit a strong topology bias, preferentially predicting links for high-degree nodes regardless of clinical validity. (D, E, G) Clinical Value Alignment via KTO. To correct this bias, the model undergoes Kahneman-Tversky Optimization (KTO). (G) Historical Phase III clinical trial outcomes serve as the ground-truth reward signal. (D) “Hard Negative” reasoning chains—structurally popular but clinically failed candidates—receive negative rewards. (E) “Novel Positive” chains—structurally sparse but clinically successful candidates—receive positive rewards. This mechanism aligns the model’s semantic reasoning with verifiable therapeutic utility rather than structural probability.

**Figure 2:**
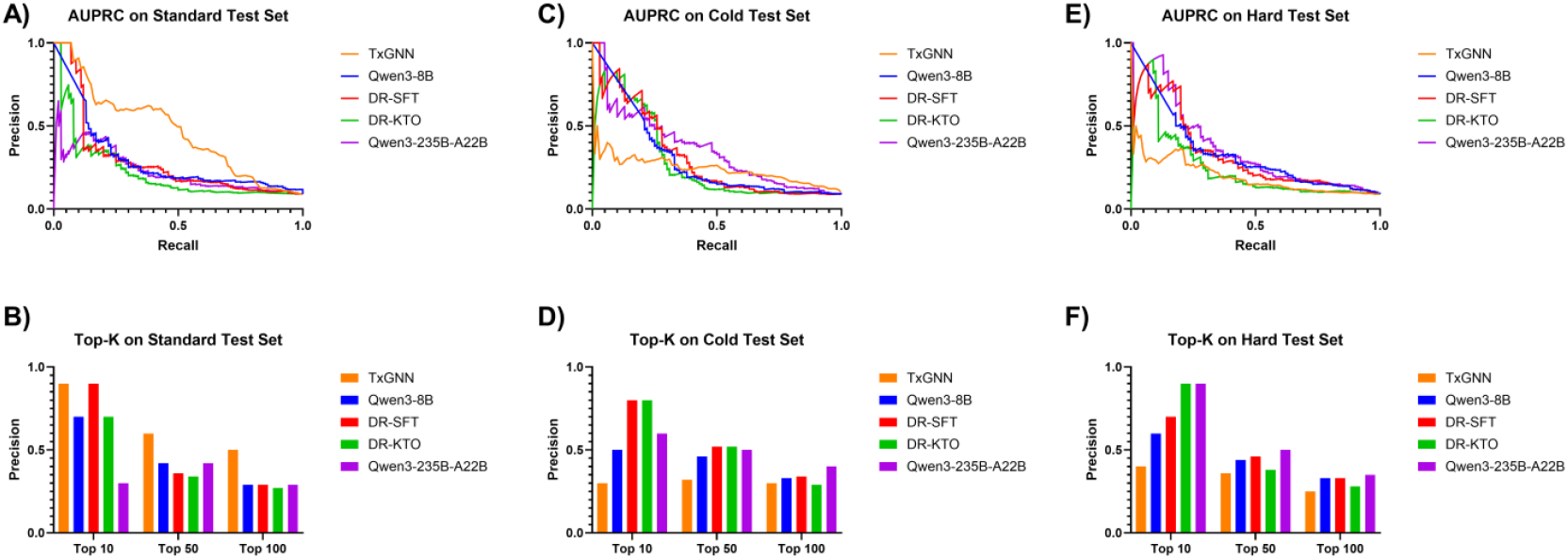
Comparative Performance of the Clinical Alignment Framework Against SOTA Baselines. Performance of the proposed framework (DR-SFT, DR-KTO) is benchmarked against the leading GNN (TxGNN) and Foundation Models (Qwen3-8B, Qwen3-235B) across three distinct 1:10 imbalanced scenarios. (A, B) Standard Setting: Precision-Recall Curves (PRC) and Top-k Precision. (C, D) Cold-Start Setting: Evaluation on novel compounds. The DR-SFT and DR-KTO models demonstrate a decisive advantage over structural baselines. Note that in Top-k metrics (D), DR-SFT slightly outperforms DR-KTO at broader retrieval ranges (*k* = 100), indicating that SFT drives the primary inductive generalization, while KTO imposes stricter selectivity. (E, F) Hard-Negative Setting: Evaluation against degree-matched negatives. Here, DR-KTO significantly outperforms DR-SFT, highlighting the necessity of outcome alignment for filtering popularity bias. Remarkably, in the critical high-confidence zone (Top-10), the lightweight DR-KTO (8B) achieves parity with the massive Qwen3-235B model, although the larger model retains an advantage in broader retrieval (*k* > 50). This underscores the efficiency of KTO in concentrating predictive density on the most clinically viable candidates.

**Figure 3:**
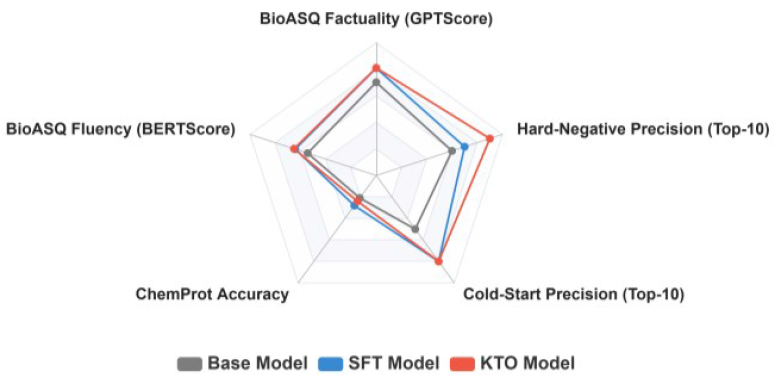
Holistic Evaluation of Biomedical Capabilities. Radar chart comparing model performance across key metrics on the Mi-RAGE, Chemprot and BioASQ dataset. The DR-SFT demonstrates the highest classification proficiency, significantly improving over the Base model, highlighting the efficacy of KG-grounded instruction tuning for mechanistic reasoning, while the KTO-aligned model (Qwen3-8B-KTO) achieves superior scores in semantic relevance metrics (GPTScore, BERTScore). It illustrates the complementary roles of semantic fine-tuning and clinical preference alignment.

### Semantic Reasoning (SFT) Solves the Cold-Start Problem

We evaluated the DR-SFT model, which utilizes Chain-of-Thought (CoT) reasoning grounded in KG paths. Unlike GNNs that rely on pre-existing edges, DR-SFT leverages the semantic understanding of drug mechanisms to bridge topological gaps. In the Cold-Start setting where GNNs collapsed, DR-SFT achieved a remarkable Top-10 Precision of 0.80 (Figure 2D), representing a substantial improvement over the graph baseline (TxGNN: ∼0.30) and the base Qwen3-8B model (0.50). This confirms that instruction tuning (SFT) is the primary driver for inductive generalization, enabling the model to infer interactions for novel compounds based purely on semantic descriptions. Furthermore, DR-SFT maintained high consistency across Standard (Figure 2B) and Cold-Start settings, demonstrating superior generalization compared to the volatile performance of graph baselines. Notably, while the KTO-aligned model also performed well, DR-SFT exhibited a slight advantage in broader retrieval metrics (Top-100), suggesting it retains a more expansive exploratory capability essential for early-stage screening.

### Clinical Alignment (KTO) Masters the Hardest Negatives

We assessed the impact of KTO-based alignment on the model’s discriminative power in high-noise environments. While DR-KTO performed comparably to DR-SFT in cold-start scenarios, it demonstrated its distinct value in the challenging Hard-Negative setting. In this scenario dominated by structurally popular but clinically ineffective decoys, DR-KTO achieved a Top-10 Precision of 0.90 (Figure 2F), significantly surpassing DR-SFT (0.70) and all GNN baselines. Remarkably, in this critical high-confidence zone (T op-10), the 8B-parameter DR-KTO model matched the performance of the massive Qwen3-235B foundation model, despite being orders of magnitude smaller. Although the larger model retained an advantage in broader retrieval (Top-50/100), DR-KTO’s parity in the top ranks suggests that explicitly penalizing clinically failed hypotheses acts as an efficient “clinical gatekeeper.” This alignment effectively filters out structurally plausible but therapeutically ineffective candidates that mislead both GNNs and standard SFT models.

### Outcome Alignment Enhances General Biomedical Reasoning and Factuality

To determine whether aligning with narrow clinical trial outcomes compromises general biomedical knowledge, we evaluated our model on two external benchmarks: ChemProt (relation extraction/classification) and BioASQ (open-ended QA) (Tabel 1)^30,31^. We benchmarked our framework against a suite of state-of-the-art open models, including the DeepSeek-R1 series and the Qwen3 family (ranging from 1.7B to 235B parameters, full results are shown in Table S1 ). On BioASQ, which demands evidence-aware, free-form generation, DR-KTO achieved a remarkable GPTScore of 4.052 and a BERTScore-F1 of 0.651. This performance not only surpasses the base Qwen3-8B model by substantial margins (GPTScore: +15%; BERTScore-F1: +20%) but, strikingly, outperforms massive foundation models such as DeepSeek-R1-671B and Qwen3-235B. This suggests that optimizing for clinical trial success implicitly teaches the model to generate responses that are more fluent, factually grounded, and clinically relevant, significantly reducing the hallucination of plausible-sounding but incorrect details. Even on the rigid multiple-choice format of ChemProt, the DR-KTO (Accuracy 0.24) maintained a solid lead over the base model (+14% relative improvement), demonstrating robust knowledge retention.

### Ablation Study: The Complementary Roles of KG-enhanced SFT and KTO

To dissect the contributions of each pipeline stage, we conducted an ablation study across training checkpoints (Table 2, full results are shown in Table S2 ). SFT with KG-grounded reasoning paths provided the most significant boost in mechanistic understanding. On ChemProt, DR-SFT model demonstrated dramatic gains over the base Qwen3-8B, improving Accuracy by 38% and Weighted-F1 by 106%, proving that graph-based rationales effectively teach the model the underlying logic of drug-protein interactions. The subsequent KTO stage further refined the model’s generation quality. On BioASQ, KTO alignment pushed BERTScore-F1 from ∼0.63 ( DR-SFT) to 0.65 ( DR-KTO ). This progression illustrates a clear synergy: KG-RAG provides the mechanistic depth required for understanding, while outcome-aligned KTO refines preference consistency, ensuring outputs align with verified clinical realities. We observed a slight performance dip for the DR-KTO compared to DR-SFT on ChemProt. This trade-off is expected and often referred to as the “alignment tax”: reinforcement learning (KTO) encourages the cautious, nuanced reasoning typical of clinical experts, which can sometimes be penalized by rigid single-choice evaluation formats that favor over-confident prediction patterns.

**Table 1.** Benchmark comparison on ChemProt (MCQ) and BioASQ (open-ended QA) between open baseline models and our outcome-aligned (DR-SFT and DR-KTO ) models. Benchmark comparison on BioASQ and ChemProt. Our 8B parameter model, aligned with clinical outcomes, achieves state-of-the-art scores on BioASQ factuality metrics, outperforming models orders of magnitude larger.

| Model name | Chemprot |  |  | BioASQ |  |
| --- | --- | --- | --- | --- | --- |
|  | Accuracy | Macro-F1 | Weighted-F1 | GPT score | BERTScore-F1 |
| DeepSeek-R1-671B | 0.48 | 0.39 | 0.50 | 3.9120 | 0.4638 |
| Qwen3-8B (Base model) | 0.21 | 0.23 | 0.16 | 3.5100 | 0.5432 |
| Qwen3-235b-a22b | **0.51** | **0.39** | **0.55** | 3.7300 | 0.5565 |
| Qwen3-8b-kto | 0.24 (+14%) | 0.23 (+0%) | 0.24 (+50%) | **4.0520** (+15%) | **0.6507** (+20%) |

**Table 2.** Ablation across base, the best DR-SFT checkpoint and DR-KTO checkpoint on ChemProt and BioASQ. Ablation analysis across training stages. DR-SFT provides the primary boost in mechanistic classification (ChemProt), while KTO further refines generation quality and factuality (BioASQ).

| Model name | Chemprot |  |  | BioASQ |  |
| --- | --- | --- | --- | --- | --- |
|  | Accuracy | Macro-F1 | Weighted-F1 | GPT score | BERTScore-F1 |
| Qwen3-8b (Base model) | 0.21 | 0.23 | 0.16 | 3.5100 | 0.5432 |
| DR-SFT ckpt-3000 | **0.28** (+33%) | **0.27** (+17%) | **0.30** (+88%) | 4.0430 (+15%) | 0.6415 (+18%) |
| DR-KTO ckpt-3500 | 0.24 (+14%) | 0.23 (+0%) | 0.24 (+50%) | **4.0520** (+15%) | **0.6507** (+20%) |

### Interpretability and Robustness: The Mechanism of Alignment

To understand the internal mechanisms driving the performance of our outcome-aligned model, we conducted a dual analysis: assessing hyperparameter sensitivity to rule out stochastic artifacts, and visualizing gradient attributions to reveal shifts in the model’s attention. We employed token-level integrated gradients to visualize feature importance for identical prompt across models (Figure 4G). As shown in the attribution heatmap, both the Base (Qwen3-8B) and DR-SFT models exhibit diffuse attention patterns. Their highest gradient salience often focuses on syntactic function tokens (e.g., prepositions “in”, whitespaces) rather than semantic content, leading to generic or format-compliant but factually shallow gene rations. In stark contrast, the DR-KTO demonstrates a decisive attention shift. It places strong, concentrated attribution on key biomedical entities (e.g., the disease term “NSCLC” in the output). This entity-centric focus suggests that clinical alignment fundamentally reorganizes the model’s latent priorities: it learns to suppress superficial linguistic cues in favor of critical therapeutic targets, directly explaining the improved factual specificity observed in the BioASQ benchmark.

**Figure 4.**
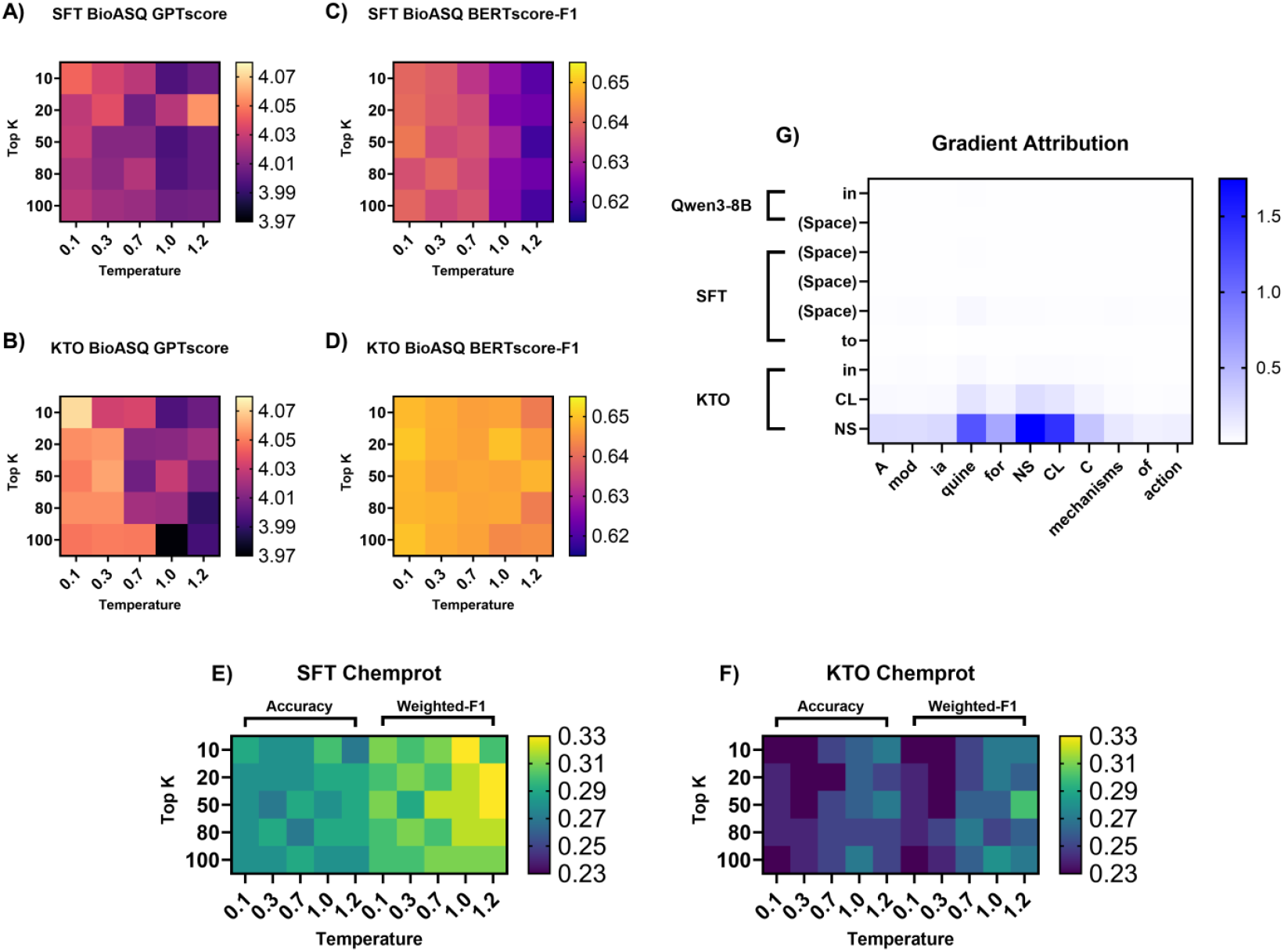
Unveiling the Mechanism of Clinical Alignment. **(A-F)** Decoding Sensitivity: Performance stability across varying Temperature and Top-K values. The KTO model shows greater sensitivity to high temperatures in open-ended generation (B subchart), suggesting that its aligned probability distribution is more peaked around clinically valid answers, necessitating lower-temperature decoding for optimal performance. Conversely, classification metrics (ChemProt) remain robust across hyperparameters for all models. **(G)** Gradient Attribution Analysis: Token-level saliency maps reveal a fundamental shift in model attention. While Base and DR-SFT (top) often focus on syntactic function words, the KTO-aligned model (bottom) concentrates high attribution scores on disease-specific entities (e.g., “NSCLC”), indicating that outcome alignment encourages evidence-grounded reasoning over superficial fluency.

We further probed the stability of these improvements by varying decoding hyperparameters (Top-K and Temperature) across ChemProt and BioASQ tasks. Overall, performance remained consistent. On ChemProt, accuracy fluctuated narrowly ( DR-SFT: 0.29–0.33; DR-KTO : 0.23–0.27), confirming that our reported gains are not artifacts of specific decoding settings. Top-K variations showed negligible impact across all tasks. On the open-ended BioASQ task, the DR-KTO exhibited higher sensitivity to temperature. While DR-SFT performance was flat, DR-KTO ‘s GPTScore degraded markedly at high temperatures

(T > 0.7). This aligns with the “cautious expert” hypothesis: preference optimization (KTO) sharpens the probability distribution around clinically valid answers. Increasing randomness (Temperature) disrupts this sharpened distribution more severely than it does the flatter distribution of the DR-SFT. This indicates that for outcome-aligned models, deterministic decoding (lower temperature) is preferable to preserve clinical precision.

### Physical Validation: Molecular Docking Confirms Binding Plausibility

To validate that the candidates generated by our framework correspond to physically viable molecules, we conducted molecular docking simulations using the DrugReAlign protocol^32^. We targeted five diverse disease areas: Type 2 Diabetes (T2DM), Non-Small Cell Lung Cancer (NSCLC), Hypertension (HT), Rheumatoid Arthritis (RA), and HIV. Figure 5 illustrates the distribution of Binding Free Energies ( Δ*G*, kcal/mol) for candidates recommended by each model. While the median affinity scores remained comparable across Base, DR-SFT, and DR-KTO—indicating all models can retrieve general drug classes—the KTO-aligned model exhibited significantly tighter score distributions. Specifically, the DR-KTO demonstrated reduced variance and a marked decrease in weak-binding outliers (long tails in the upper range of Δ*G*) across all five disease domains. For instance, in the NSCLC panel, the interquartile range (IQR) of the DR-KTO was narrower than that of the DR-SFT, with fewer candidates exceeding the -6.0 kcal/mol threshold (a proxy for weak affinity). This convergence suggests that clinical preference alignment implicitly promotes physical plausibility: by penalizing clinical failures, the model learns to filter out structurally or mechanistically mismatched compounds, thereby concentrating its recommendations on high-confidence binders. This property is critical for downstream experimental validation, as it minimizes the risk of allocating wet-lab resources to low-affinity false positives.

**Figure 5.**
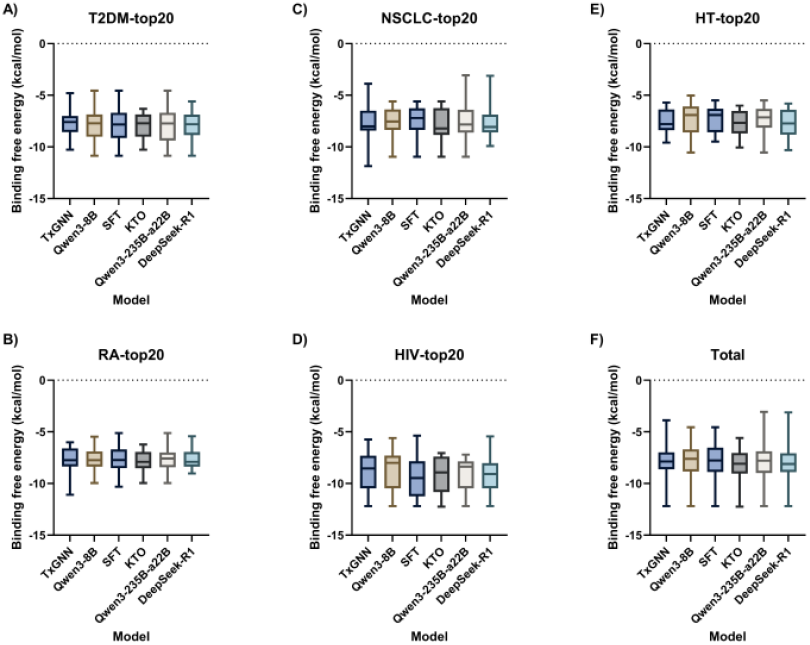
Physical Validation of Model Predictions via Molecular Docking. **(A-F)** Boxplots of binding free energies (Δ*G*) for drug candidates recommended by TxGNN, Qwen3-8B (Base), SFT, KTO, Qwen3-235B-A22B and DeepSeek-R1 models across five disease targets. While median affinities are similar, the KTO model consistently exhibits tighter distributions with fewer weak-binding outliers (high Δ*G* values) compared to SFT and Base models. This indicates that clinical alignment filters out physically implausible candidates, improving the signal-to-noise ratio for downstream screening.

**Figure 6.**
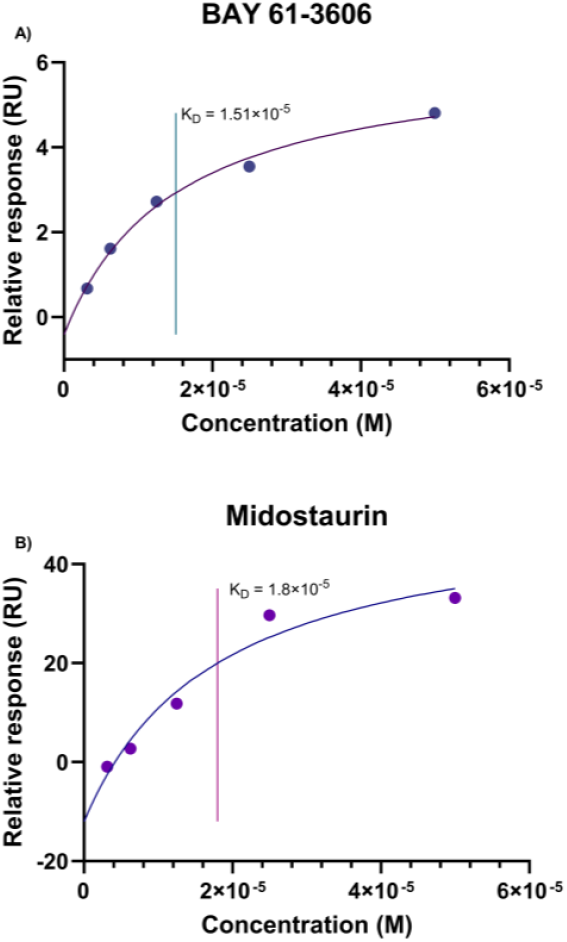
Binding affinity of BAY 61-3606 (A) and Midostaurin (B) to Flt3 protein by SPR. Steady-state binding responses at five different compound concentrations were plotted against concentration to generate dose–response curves. The experimental data points were fitted with a 1:1 binding model (solid curve) to determine the dissociation constant (KD).

### Case Study: Identification of BAY 61-3606 as a Novel FLT3 Binder for AML

For experimental validation, we continued employing the DrugReAlign pipeline to systematically prioritize candidates based on a “High-Confidence, High-Affinity” strategy. Specifically, we screened model-predicted candidates for Acute Myeloid Leukemia (AML) to identify those exhibiting exceptional binding scores in orthogonal molecular docking simulations against FLT3, a critical therapeutic target. BAY 61-3606 emerged as the standout candidate from this structural filtration. While documented as a preclinical Syk inhibitor with observed cytotoxicity in leukemia cells^33^, its direct interaction with FLT3 remains unreported, satisfying our criteria for both mechanistic novelty (unknown target) and translational feasibility (known phenotype). This dual profile—strong predicted affinity versus unexplored specific mechanism—positions BAY 61-3606 as an optimal test case to verify the model’s capacity for mechanism deconvolution.

**Table 3:**
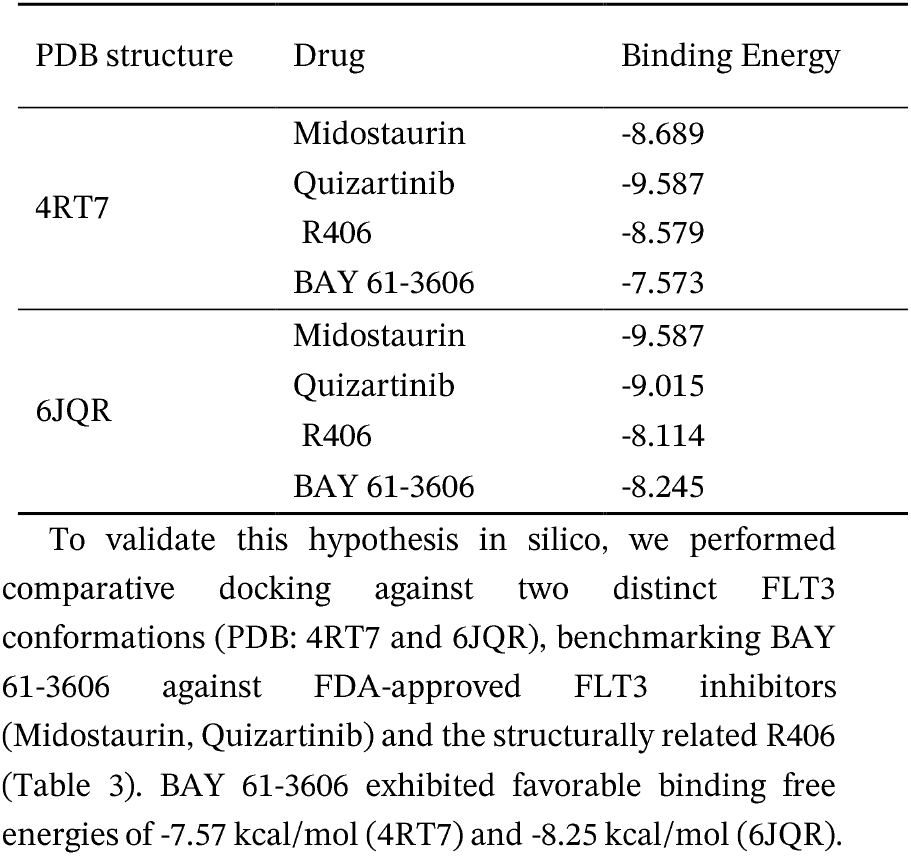
Comparative Docking Scores (In Silico Validation)

| PDB structure | Drug | Binding Energy |
| --- | --- | --- |
| 4RT7 | Midostaurin | -8.689 |
|  | Quizartinib | -9.587 |
|  | R406 | -8.579 |
|  | BAY 61-3606 | -7.573 |
| 6JQR | Midostaurin | -9.587 |
|  | Quizartinib | -9.015 |
|  | R406 | -8.114 |
|  | BAY 61-3606 | -8.245 |

Crucially, in the 6JQR conformation, its affinity surpassed that of R406 ( -8.11 kcal/mol), a compound with known anti-leukemic effects linked to the FLT3 pathway. Although weaker than nanomolar-potency inhibitors like Quizartinib (-9.02 kcal/mol), these scores fall well within the range of biologically active hits, suggesting BAY 61-3606 possesses non-trivial binding potential for FLT3.

To experimentally confirm the prediction, we conducted SPR binding assays with purified FLT3 protein for both BAY 61-3606 and midostaurin under matched conditions. While the absolute values of K_D_ obtained from SPR measurements may be affected by experimental pipelines, the results provide a robust basis for relative comparison. Notably, BAY 61-3606 demonstrated FLT3 binding at a level comparable to that of midostaurin, a clinically validated FLT3 inhibitor. This finding substantiates our computational predictions and further supports the hypothesis that BAY 61-3606 may act as a direct FLT3 binder, warranting further functional investigation in the context of AML.

## Discussion

### Shifting from Structural Probability to Clinical Utility

The central premise of this work is that effective drug repurposing requires a paradigm shift from “structural probability”—the optimization objective of traditional GNNs—toward “clinical utility” grounded in verifiable outcome data. Our study validates this hypothesis through a hierarchical evaluation strategy that progresses from macroscopic statistical inference to microscopic biophysical verification. At the broadest level, results on the MiRAGE benchmark reveal that while GNNs excel in transductive settings, they fail catastrophically in real-world “Cold-Start” and “Hard-Negative” scenarios. In contrast, our KG-augmented DR-SFT model successfully bridges these topological gaps via semantic reasoning. Crucially, at the deepest level of validation, the DR-KTO alignment mechanism proved essential for filtering popularity bias, a capability orthogonally confirmed by the discovery and experimental validation (via SPR) of BAY 61-3606 as a novel high-affinity FLT3 binder. Collectively, these findings demonstrate that integrating semantic generalization with value-based alignment offers a robust framework that transcends the limitations of purely topological approaches, delivering candidates that are not only structurally plausible but clinically viable.

### Semantic Reasoning and Early-Stage Strategy (SFT)

A critical bottleneck in traditional repurposing is the “Cold-Start” problem: predicting indications for novel compounds with no prior graph connectivity. DR-SFT’s achievement of 0.80 Top-10 Precision—far surpassing the best GNN baseline (TxGNN, 0.30)—signals a fundamental shift from neighbor propagation to mechanistic reasoning. This is corroborated by our ChemProt ablation studies, where learning the semantic logic of interactions allowed the model to generalize to unseen entities based on literature descriptions alone. Consequently, we identify DR-SFT as the optimal engine for the “Early Screening” phase. In scenarios involving new chemical entities (NCEs) or emerging pathogens where structural priors are absent, SFT should be deployed to maximize recall and explore chemical space beyond the training graph, serving as the primary inductive generator before graph-based methods can even operate.

### Clinical Alignment and Late-Stage Strategy (KTO)

While SFT expands the search space, this work validates KTO as a critical “clinical gatekeeper.” In the Degree-Matched setting, GNNs were misled by “hub” nodes—popular but ineffective drugs—whereas the KTO-aligned model achieved 0.90 Precision, effectively filtering out these “scientific bubbles.” Gradient attribution maps reveal that KTO shifts attention from superficial syntactic cues to specific disease subtypes, proving that penalizing clinical failures teaches the model to distinguish associative popularity from causal efficacy. Thus, DR-KTO is uniquely suited for the “Lead Optimization” and “Portfolio Review” stages. Once a candidate pool is established, KTO should be employed to rigorously filter false positives in dense, well studied therapeutic areas (the “Hard Negative” regime). Furthermore, for tasks requiring mechanistic interpretation—such as generating regulatory reports—KTO is the preferred engine due to its enhanced factual grounding on benchmarks like BioASQ.

### Physical Reality and Translational Potential

Ultimately, computational prediction must correspond with physical reality. Our molecular docking simulations demonstrate that the model’s “clinical intuition” is grounded in biophysical plausibility, consistently recommending candidates with tighter bindi ng energy distributions. The identification of BAY 61-3606 as a high-affinity FLT3 binder serves as a quintessential example. Despite the lack of direct training edges, the model inferred this connection via pathway crosstalk, a hypothesis subsequently confirmed by our SPR assays. Since BAY 61-3606 is an investigational small molecule, this newly predicted binding suggests tangible potential for commercial development in AML indications. This end-to-end validation establishes the “SFT-for-Exploration, KTO-for-Validation” paradigm not merely as a ranking algorithm, but as a reliable engine for nominating physically viable candidates for preclinical trials.

### Limitations and Future Directions

Despite these advances, two critical limitations remain. First, while DR-KTO instills a form of “clinical judgment,” the biological essence of this judgment remains opaque. This opacity is twofold: it stems not only from the “black box” nature of LLMs but from the inherent complexity of pharmacology itself—where the precise determinants of Phase III success (e.g., subtle pharmacokinetic properties or off-target toxicities) are often undefined even by human experts. The model likely captures high-dimensional heuristic patterns that correlate with success, but fully explicating these rules remains an open challenge. Second, the system is designed to augment, not replace, human expertise. Our results show that while the model excels at triage, its outputs are probabilistic and require rigorous verification by pharmacologists. Furthermore, the model’s performance is sensitive to prompt engineering; domain experts unaware of prompt nuances may inadvertently elicit suboptimal responses. Future work must focus on developing intuitive interfaces that bridge the gap between AI capabilities and pharmacological workflows, ensuring that human experts remain the final arbiters of therapeutic decision-making.

### Conclusion

This study addresses the critical limitations of topological biases in traditional drug repurposing by introducing a unified framework that couples semantic reasoning with clinical outcome alignment. Our results demonstrate that while GNNs falter in inductive and high-noise scenarios, our KG-augmented SFT model effectively overcomes the “Cold-Start” barrier for novel compounds, while KTO alignment acts as a robust clinical gatekeeper, suppressing the high-degree false positives that plague standard algorithms. Validated by rigorous benchmarking, molecular docking, and the experimental confirmation of BAY 61-3606 as a novel FLT3 binder, our framework bridges the gap between computational inference and physical reality. Ultimately, this work establishes a new paradigm for drug discovery—moving from structure-based probability to outcome-driven utility—offering a scalable, evidence-grounded path to accelerate the identification of clinically viable therapeutics.

## Acknowledgements

This work was supported by the Science and Technology Development Fund of Macau SAR (No.: 0002/2025/NRP, 0049/2024/AGJ), the University of Macau (No.: MYRG-CRG2023-00007-ICMS-IAS, MYRG-GRG2024-00268-ICMS -UMDF, and SHMDF-AI/2026/003 with Dr. Stanley Ho Medical Development Foundation), the Prevention and Control of Emerging and Major Infectious Diseases-National Science and Technology Major Project(2025ZD01907900), and High-performance Computing Platform of Peking University.

We thank Prof. Qi Wang and Prof. Wusheng Xiao and their groups for the convenience of wet lab using during the project.

We acknowledge the use of GitHub Copilot ([GitHub Copilot · Your AI pair programmer](https://github.com/features/copilot), powered by OpenAI’s Codex) during the preparation of this manuscript. The tool was used exclusively for code debugging, syntax optimization, language refinement and polishing English grammar. The authors reviewed and verified all AI-generated content and took full responsibility for the accuracy and integrity of the final publication.

This paper was typeset with the bioRxiv word template by @Chrelli: www.github.com/chrelli/bioRxiv-word-template

## Author contributions

Ruihu D U: Conceptualization, Methodology, Software, Data curation, Formal analysis, Investigation, Writing – original draft, Visualization. Mancheong FUNG : Conceptualization, Writing – review & editing. Yuanjia Hu : Conceptualization, Resources, Funding acquisition, Project administration, Supervision, Writing – review & editing. Dongyang Liu : Resources, Funding acquisition, Project administration, Supervision, Writing – review & editing.

## Data availability

All data and code are available, please refer to <https://github.com/cat-on-tree/drkto/> for more information.

## Materials and Methods

### Data Resources and Preparation

To support mechanistic reasoning and outcome alignment, we integrated three complementary resources:

1. PrimeKG^6^: A knowledge graph containing 129,375 nodes and >4 million relations. It serves as the retrieval substrate for our KG-RAG framework.
2. RepoDB^34^: A clinical benchmark comprising 6,677 approved and 4,123 failed drug-indication pairs, providing the large-scale positive and negative signals required for both SFT and KTO stages.
3. DrugRepoBank^35^: A manually curated resource of drug repurposing associations enriched with DrugBank metadata, serving as the seed corpus for high-quality inductive reasoning.

To ensure semantic consistency, all entities across datasets were standardized to a unified vocabulary (DrugBank/ChEMBL for drugs, UMLS/MeSH for diseases), facilitating seamless linking between textual instructions and the PrimeKG structure. We constructed a hierarchical training corpus through a multi-stage pipeline (illustrated in Figure S1):

We transformed DrugRepoBank entries into multi-turn inductive dialogues. Each dialogue consists of five reasoning rounds, guiding the model to sequentially analyze: (1) Drug Mechanism, (2) Candidate Diseases, (3) Potential Pathways, (4) Evidence Correlation, and (5) Final Recommendation. This protocol explicitly trains the model to attribute evidence before concluding.

To scale up the training data, we processed RepoDB entries into bidirectional QA pairs. Crucially, raw RepoDB pairs lack reasoning details. We therefore passed all RepoDB pairs through our KG-RAG adaptation pipeline (described in **KG-RAG** section) to generate synthetic CoT rationales rooted in PrimeKG paths. The samples generated underwent self-consistency checks. Only samples with logically consistent reasoning traces were retained.

For SFT, we combined the 5-round dialogues from DrugRepoBank with the consistency-filtered positive samples (Approved) from the augmented RepoDB. This hybrid dataset ensures the model learns both deep interactive reasoning (from DrugRepoBank) and broad knowledge coverage (from RepoDB) ; For KTO, we utilized the full spectrum of the augmented RepoDB data, including both Approved (Success) and Failed pairs. The inclusion of failed trials—augmented with their corresponding (plausible but ultimately incorrect) reasoning traces—is critical for teaching the model to discriminate between mechanistically sound but clinically invalid hypotheses.

### Reasoning-Augmented Data Generation (KG-RAG)

We adapted a KG-RAG pipeline, this module is directly employed from Medreason to generate synthetic Chain-of-Thought (CoT) rationales for the RepoDB dataset, enriching clinical pairs with mechanistic evidence^36^. First was the entity linking and path retrieval. Given a query-answer pair (𝒬, 𝒜), we first extracted biological entities and linked them to the PrimeKG knowledge graph. We utilized a hybrid resolver that combines exact matching with embedding-based similarity (threshold *τ* = 0.85). For each linked pair 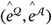, we enumerated candidate shortest paths 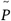 within a maximum hop distance *d*_*max*_.

Then there were the semantic pruning and aggregation. To filter out irrelevant topological connections, candidate paths were re-ranked by a filtering LLM conditioned on the specific repurposing context. We retained the top-*K* paths (empirically *K* = 3) to form the aggregated evidence set Π.

Finally, CoT synthesis and quality control were performed. A generator LLM synthesized a reasoning trace 𝒞 based on the retrieval evidence Π :

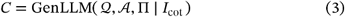

This ensured the logical validity of the generated rationales; we also applied a self-consistency check: an independent evaluator model verified whether the generated reasoning 𝒞 logically supported the ground-truth target 𝒜. Only samples passing this verification were retained for the subsequent training phases.

For more details, please refer to the original article^36^.

### Supervised Fine-Tuning (SFT)

The SFT stage aims to instill mechanistic reasoning capabilities and domain knowledge into the backbone model. Based on Qwen3-8B^37^, we utilized the composite dataset 𝒟_SFT_, which comprises high-quality 5-round inductive dialogues from DrugRepoBank and consistency-filtered approved samples from RepoDB (as **Data Resources and Preparation** illustrated ).

Let *x* denote the input instruction (e.g., the repurposing query) and *y* denote the target output sequence, which consists of the reasoning trace followed by the final recommendation. The model parameter *θ* is optimized using the standard autoregressive negative log-likelihood (NLL) ob-jective:

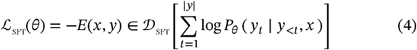

We fine-tuned the model for 3 epochs. The checkpoint achieving the lowest validation NLL was selected as the reference model *π* _ref_ (or *θ*_0_ ) to initialize the subsequent alignment stage.

### Clinical Outcome Alignment via KTO

To align the model’s latent preferences with real-world clinical utility, we employed KTO^25^. Unlike PPO or DPO which require paired preference data (Winner vs. Loser), KTO is designed for unpaired binary signals, making it ideal for leveraging clinical trial data where “Success” (Approved) and “Failure” (Terminated/Withdrawn) are distinct, independent events.

We utilized the 𝒟 _KTO_ dataset, which includes both approved ( *o* = 1) and failed ( *o* = 0) drug-indication pairs from RepoDB, along with their generated reasoning traces.

#### Implicit Reward Definition

We define the implicit reward ratio *r* _*θ*_ *(x, y)* as the log-likelihood difference between the current policy *π*_*θ*_ and the frozen SFT reference policy *π* _ref_:

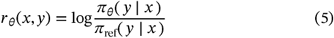

#### Objective Function

The KTO objective treats clinical outcomes as prospect-theoretic rewards. We maximize the likelihood of clinically successful generations relative to the reference model while suppressing unsuccessful ones. The loss function is formulated as a binary classification objective over the implicit reward:

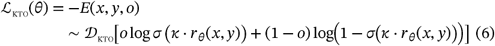

where *σ* is the sigmoid function, and *κ* is a hyperparameter controlling the steepness of the preference curve (i.e., the strength of the alignment signal). This objective effectively pushes the probability mass of the model towards clinically validated mechanisms and away from hypotheses associated with trial failures, acting as a “soft” reinforcement learning signal without the need for an explicit reward model.

The pseudo-code for the three steps is listed in Supplementary Note 1.

## Evaluation

### Drug Repurposing Benchmarks

To rigorously assess repurposing performance, we utilized the MiRAGE^26^ benchmark derived from CTD^27^. We benchmarked our framework against four state-of-the-art GNNs: RGCN, HGT, HAN, and TxGNN^9^, implemented via PyTorch Geometric^28,29^. The GNN models were trained on the PrimeKG graph (80:20 train-validation split) with type-constrained negative sampling. Crucially, node features were randomly initialized to isolate topological learning capability from semantic priors. For the test set, we aligned MiRAGE data with PrimeKG; entries involving entities absent from the graph (42.2%) were excluded from GNN evaluation to ensure topological reachability. We constructed three distinct testing scenarios under fixed random seeds: Standard, Cold-Start (defined as disease nodes with degree < 3 in the training graph), and Degree-Matched (Hard Negatives). Evaluation was performed under both balanced (1:1) and realistic imbalanced (1:10) negative sampling ratios. For LLM s, we evaluated Qwen3-8B (Base), our SFT and KTO variants, and the large-scale Qwen3-235B-A22B reference. To ensure a unified metric space, we adopted a logit-based scoring approach: models were prompted to classify associations (e.g., “Will [Drug] treat [Disease]? Answer True or False”), and the normalized log-probabilities of the target tokens were extracted to derive continuous confidence scores for AUPR C and Top-10 Precision. Detailed hyperparameters and prompts are provided in the GitHub repository.

#### General Biomedicine task Benchmarks

To assess the broader biomedical reasoning capabilities of our model beyond specific repurposing tasks, we evaluated performance on two complementary resources: ChemProt^30^ and BioASQ^31^. ChemProt was formatted as a multiple-choice QA task, this benchmark evaluates the model’s ability to classify chemical-protein interactions grounded in knowledge graph evidence. We report Accuracy, Macro-F1, and Weighted-F1. BioASQ was adapted as an open-ended QA task, this benchmark tests factuality and reasoning breadth. Generated responses were scored against gold-standard references using GPTScore, BERTScore-F1, and ROUGE. We compared our SFT and KTO models against a comprehensive suite of open-source foundation models, including the Qwen3 series (ranging from 1.7B to the massive 235B-A22B MoE) and the DeepSeek-R1 distilled series (1.5B to 70B). All baselines were evaluated using identical prompt templates and scoring protocols to ensure fair comparison.

#### Hyperparameter Sensitivity and Interpretability

Following the technical report of the Qwen3 series, we conducted a hyperparameter sensitivity analysis by systematically varying top_k and temperature settings during inference to determine their impact on generation quality. Furthermore, to elucidate the “black box” mechanisms of our alignment strategy, we performed gradient-based attribution analysis. By visualizing input-output attention patterns, we investigated how the focus of the model shifts across the Base, SFT, and KTO stages, specifically examining whether clinical alignment drives the model to attend more heavily to mechanistic entities versus generic tokens.

#### Orthogonal Physical Validation via Molecular Docking

To verify that our model’s semantic predictions correspond to physical reality, we employed DrugReAlign, a docking-based evaluation protocol^32^. We adopted a disease-centric comparative setup to pit our LLM directly against the strongest GNN baseline (TxGNN):

1. Target Selection: For a selected disease, we first retrieved the top-N (N = 20) drug candidates predicted by TxGNN and identified their primary protein targets.
2. Quota-Matched Recommendation: We then prompted LLM (using multi-source context) to recommend an equal number of candidates for the same targets, ensuring a strict head-to-head comparison controlling target difficulty.
3. Binding Energy Assessment: All recommended candidates (from both TxGNN and LLM) were docked against their respective protein structures using a standardized pipeline (identical preparation and scoring 1. functions). We compared the distribution of binding free energies (kcal/mol) to quantify molecular plausibility, operating on the premise that a clinically valid repurposing model should recommend drugs with thermodynamically favorable binding affinities.

#### Surface Plasmon Resonance (SPR) Assays

1. Protein Immobilization. The binding affinity between the target protein FLT3 and candidate small molecules was measured using a Biacore 8K^+^ instrument (Cytiva, Washington, USA). Recombinant human FLT3 protein with a His-tag (Elabscience Biotechnology Co., Ltd, Wuhan, China) was reconstituted in ultra-pure water to a concentration of 1 mg/m L. The protein was then diluted in 10 mM sodium acetate buffer (pH 4.5) and immobilized onto the sensor chip surface via standard amine coupling using EDC/NHS (1-ethyl-3-(3-dimethylaminopropyl) carbodiimide hydrochloride / N-hydroxy succinimide) chemistry. Following immobilization, unreacted ester groups were blocked with 1 M ethanolamine-HCl (pH 8.5). To minimize non-specific binding, the surface was conditioned with 10 mg/mL BSA.
2. Running Buffer Preparation. The buffer (1 L) consisted of 20 mM Tris, 300 mM NaCl, 10 mM MgCl _2_· 6H_2_O, and 50 μL Tween-20 dissolved in ultra-pure water and filtered through a 0.22 μm membrane. For solvent correction and analyte dilution, a stock solution (Liquid A) was prepared by taking 50 mL of the base buffer. The final running buffer (containing 5% DMSO to ensure compound solubility) was prepared by mixing 950 mL of base buffer with 50 mL of DMSO.
3. Analyte Preparation and Dilution. Two compounds were tested: Midostaurin (positive control) and BAY 61-3606 (test candidate). Both were dissolved in DMSO to prepare 10 mM stock solutions.
  3.1 Intermediate Dilution: For Midostaurin, 3 μL of stock was mixed with 12 μL DMSO; for BAY 61-3606, 1.5 μL of stock was mixed with 13.5 *μ*L DMSO. Both mixtures were subsequently diluted with 285 μL of Liquid A and centrifuged at 12,000 rpm for 5 minutes to remove potential aggregates.
  3.2 Serial Dilution: A 240 μL aliquot of the supernatant was taken for each compound. Ten-point serial dilutions (1:2) were prepared using the running buffer (containing 5% DMSO) to generate a concentration gradient. The final injection volume for each cycle was 120 μL.
  3.3 Solvent Correction. To compensate for bulk refractive index changes caused by DMSO, a solvent correction curve was generated ranging from 4.5% to 5.8% DMSO. Specifically, calibration solutions were prepared by mixing varying ratios of a 4.5% DMSO buffer (9 .5 mL Liquid A + 450 μL DMSO) and a 5.8% DMSO buffer (9.5 mL Liquid A + 580 μL DMSO). Four representative points (Points 1, 2, 7, and 8) were used for the calibration curve, with 120 μL injected per cycle. A 50% DMSO aqueous solution served as the normalization solution.
4. Data Acquisition and Analysis. Binding kinetics were monitored at 25° C. The association and dissociation phases were recorded in real-time. Raw sensorgrams were solvent-corrected, reference-subtracted, and analyzed using the Biacore Insight Evaluation Software. Equilibrium dissociation constants ( K _D_) were derived by fitting the data to a 1:1 Langmuir binding model or a steady-state affinity model depending on the kinetic profile.

**Figure S1:**
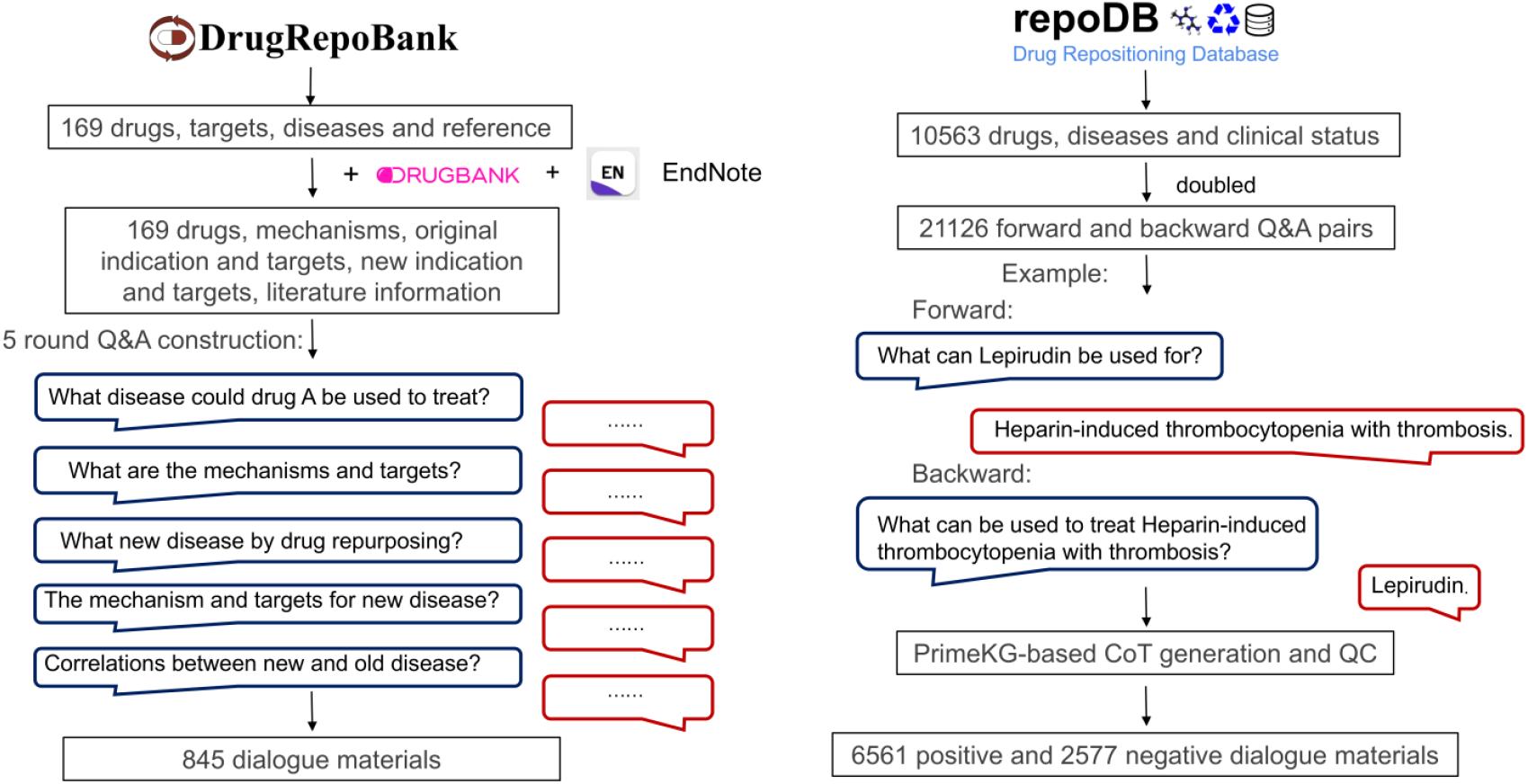
Data Processing Pipeline. The pipeline integrates heterogeneous resources (DrugRepoBank, RepoDB, PrimeKG) into task-specific training corpora via a multi-stage augmentation process. **(Left)** High-quality entries from DrugRepoBank are structured into 5-round inductive dialogues, explicitly modeling the sequential logic of mechanism analysis, pathway correlation, and recommendation; **(Right)** Clinical pairs from RepoDB (both Approved and Failed) are formatted as bidirectional QA pairs and enriched via the KG-RAG framework. Synthetic Chain-of-Thought (CoT) rationales are generated based on PrimeKG paths and filtered through a self-consistency quality control (QC) check.

**Figure S2:**
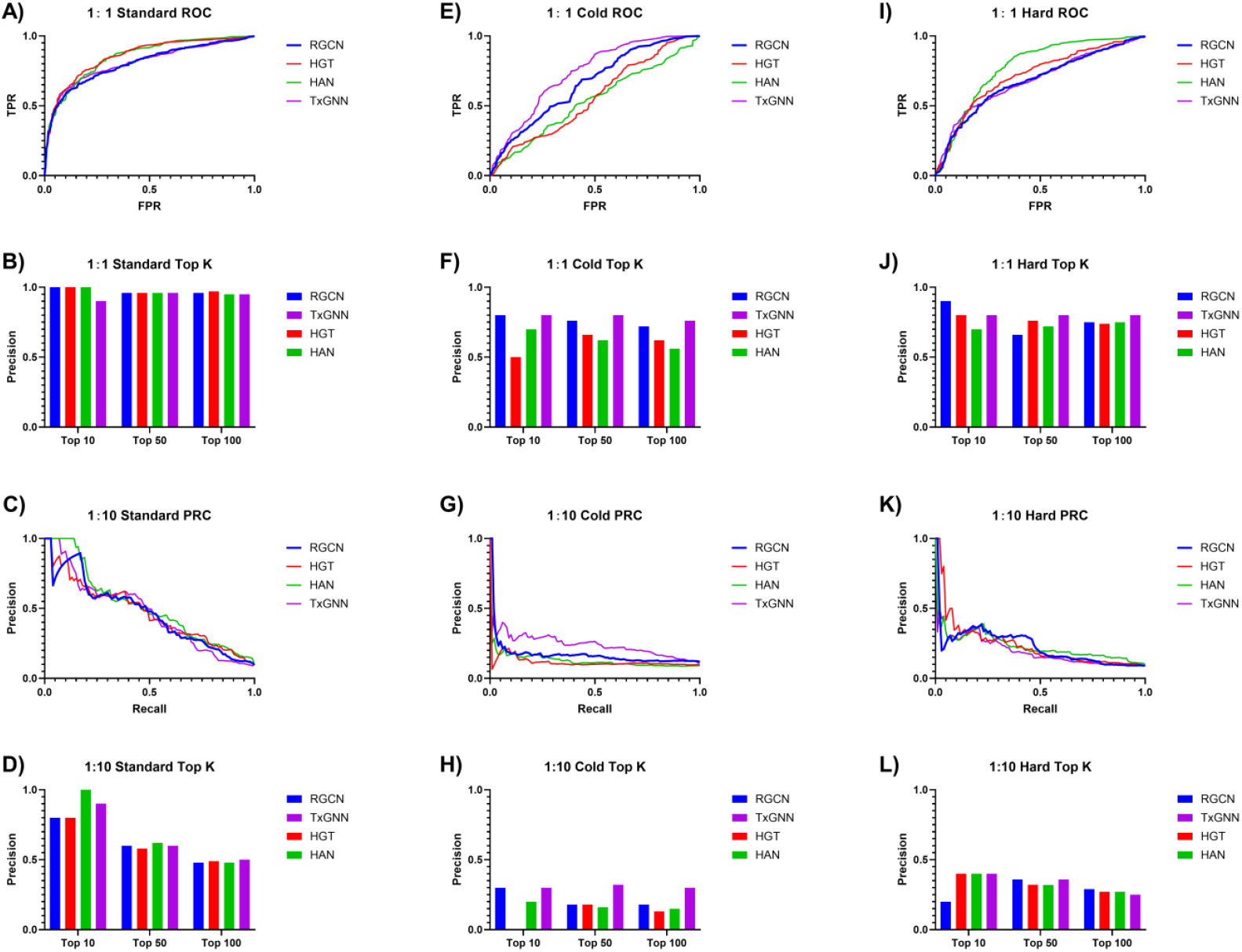
Performance Benchmarking of Graph Neural Network Baselines Across Balanced and Imbalanced Scenarios. This figure evaluates the robustness of GNN architectures (RGCN, HGT, HAN, TxGNN). The panels are organized vertically by scenario: (A–D) Standard Setting, (E– H) Cold-Start Setting, and (I–L) Degree-Matched (Hard-Negative) Setting. Within each column, the rows represent: 1:1 ROC (1st row), 1:1 Top-K Precision (2nd row), 1:10 PRC (3rd row), and 1:10 Top-K Precision (4th row). Observations: In the Standard setting (A–D), GNNs demonstrate robust performance regardless of data balance. Notably, in the challenging Cold-Start (E, F) and Hard-Negative (I, J) scenarios, GNNs maintain acceptable predictive capability under balanced (1:1) conditions. However, a catastrophic performance collapse is observed specifically in the realistic, highly imbalanced (1:10) settings (G, H for Cold-Start; K, L for Hard-Negative). This indicates that while GNNs can handle structural difficulties or class imbalance individually, they fail to generalize when simultaneously confronted with both topological sparsity/bias and realistic negative-sample dominance.

**Table S1.**
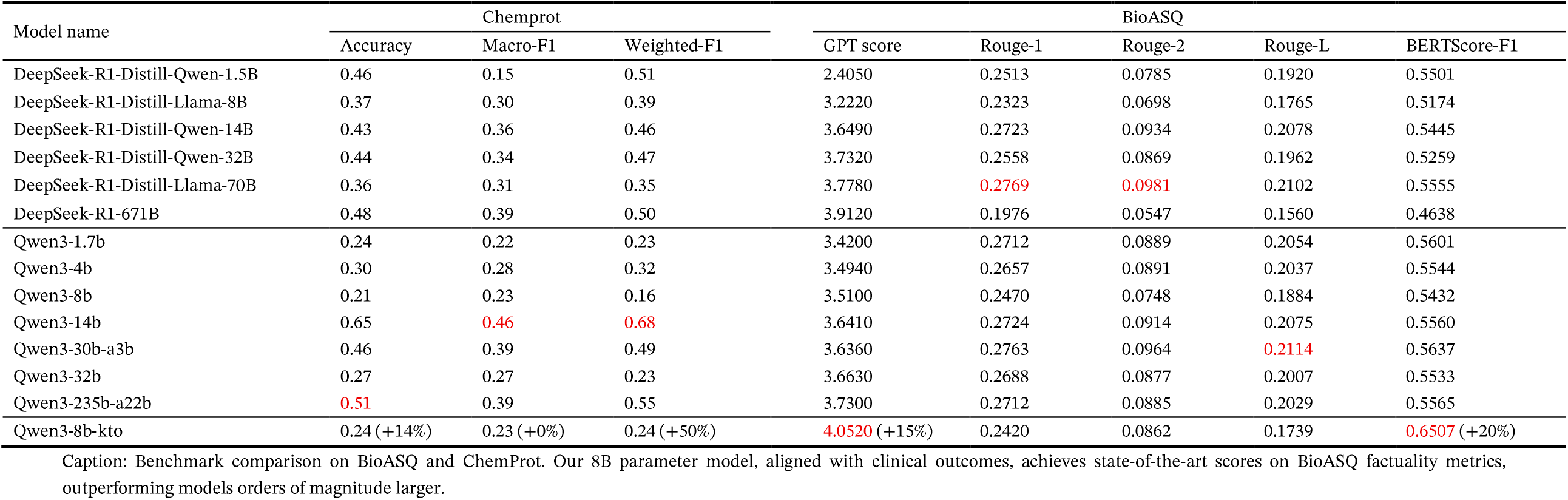
Benchmark comparison on ChemProt (MCQ) and BioASQ (open-ended QA) between open baseline models and our outcome-aligned (SFT and KTO) model.

**Table S2.**
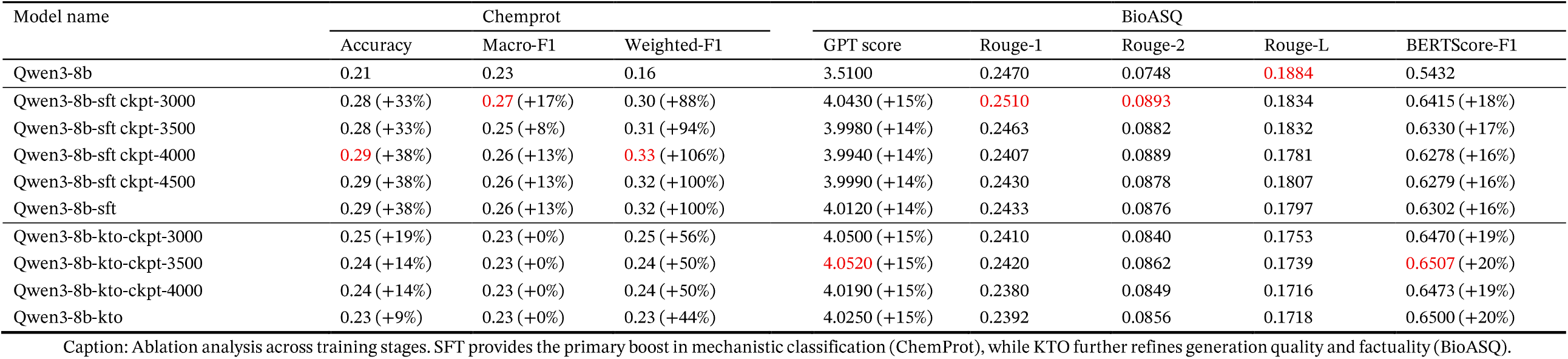
Ablation across base, SFT checkpoints, and KTO checkpoints on ChemProt and BioASQ.

## Supplementary Note 1: Outcome-Aligned KG-RAG Repurposing Framework

~~~
Input: Raw corpora D0, Knowledge Graph G
Output: Outcome-aligned model parameters θ
// Stage 1: Reasoning-Augmented Data Generation
Initialize D_reason ← ∅
for each (Q, A) in D0 do
  E_Q, E_A ← EntityLinking(Q, A, G)
  Π ← PathRetrieval(G, E_Q, E_A)
  Π_ranked ← SemanticPruning(Π, Q, A)
  C ← GenerateCoT(Q, A, Π_ranked)
  if SelfConsistencyCheck(C, A) is True then
      D_reason ← D_reason ∪ {(Q, C, A)}
  end if
end for
// Stage 2: Supervised Fine-Tuning (SFT)
θ ← InitializeBackbone()
D_SFT ← FilterSuccessful(D_reason) // Only use approved drugs for SFT
θ_ref ← TrainSFT(θ, D_SFT, Objective=NLL)
// Stage 3: Clinical Alignment (KTO)
// Note: KTO is typically offline. We use pre-generated outputs y with labels o.
D_KTO ← {(Q, y, o)} // y is model generation, o is clinical label (1=Success, 0=Fail)
θ ← θ_ref
for epoch = 1 to E_KTO do
  for batch B ⊂ D_KTO do
      L_batch ← 0
      for (Q, y, o) in B do
          // Compute log-probabilities under current policy and reference
          log_p_θ ← Forward(θ, Q, y) log_p_ref ← Forward(θ_ref, Q, y)
          // Likelihood ratio
          log_rho ← log_p_θ-log_p_ref
          // Sigmoid confidence transform
          p ← Sigmoid(κ * log_rho)
         // Binary preference loss (maximize p if o=1, minimize if o=0)
         L_item ←-[ o * log(p) + (1-o) * log(1-p) ]
         L_batch ← L_batch + L_item
      end for
      θ ← Update(θ, ∇ L_batch)
  end for
end for
return θ
~~~

