## Supplementary Materials for "Overcoming Topology Bias and Cold-Start Limitations in Drug Repurposing: A Clinical-Outcome-Aligned LLM Framework"

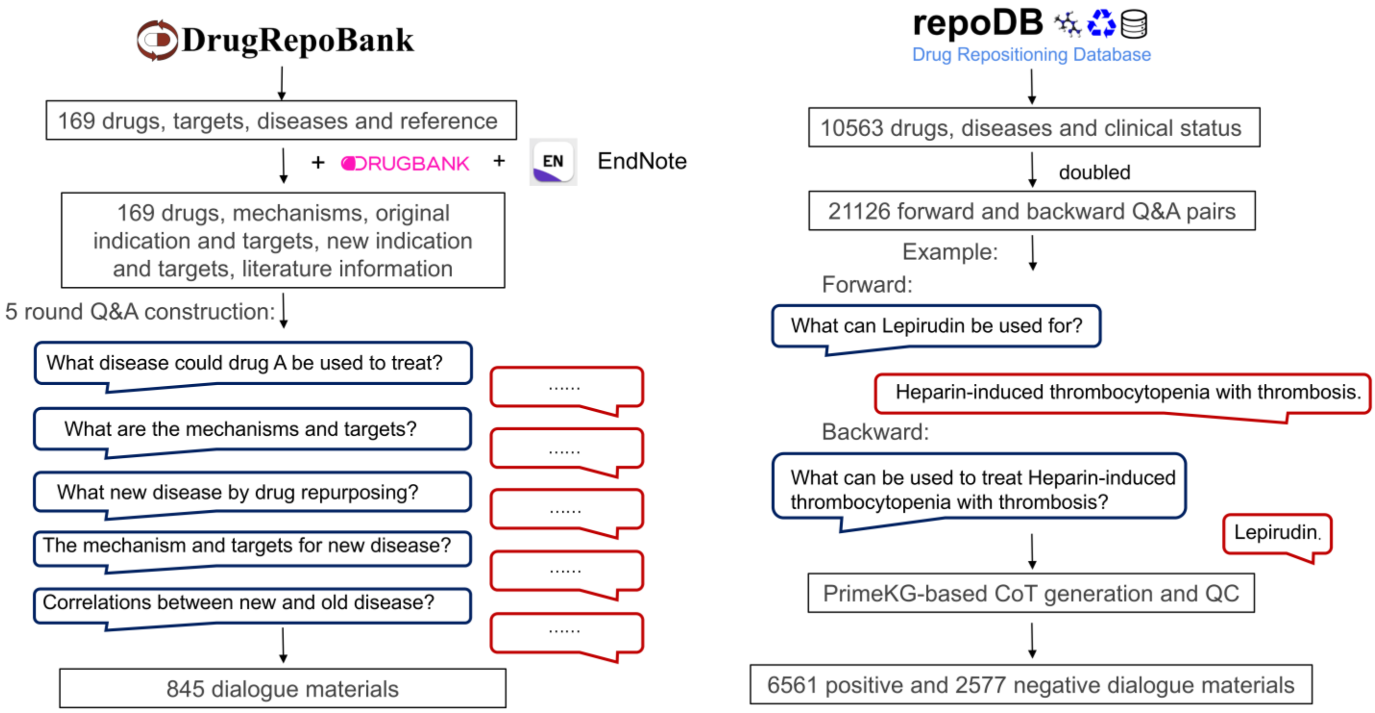


**Figure S1: Data Processing Pipeline.** The pipeline integrates heterogeneous resources (DrugRepoBank, RepoDB, PrimeKG) into task-specific training corpora via a multi-stage augmentation process. **(Left)** High-quality entries from DrugRepoBank are structured into 5-round inductive dialogues, explicitly modeling the sequential logic of mechanism analysis, pathway correlation, and recommendation; **(Right)** Clinical pairs from RepoDB (both Approved and Failed) are formatted as bidirectional QA pairs and enriched via the KG-RAG framework. Synthetic Chain-of-Thought (CoT) rationales are generated based on PrimeKG paths and filtered through a self-consistency quality control (QC) check.

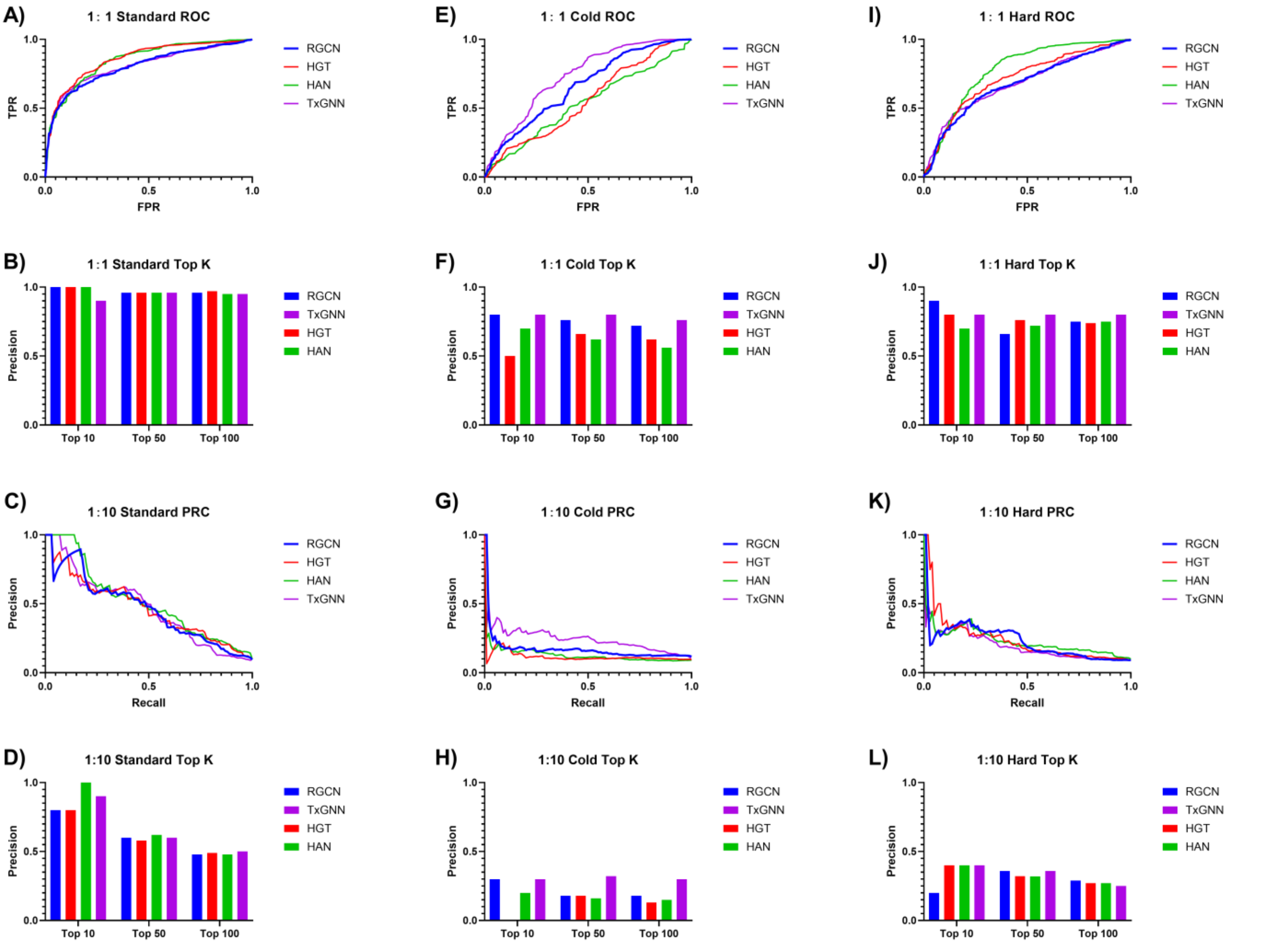


**Figure S2: Performance Benchmarking of Graph Neural Network Baselines Across Balanced and Imbalanced Scenarios.** This figure evaluates the robustness of GNN architectures (RGCN, HGT, HAN, TxGNN). The panels are organized vertically by scenario: (A–D) Standard Setting, (E–H) Cold-Start Setting, and (I–L) Degree-Matched (Hard-Negative) Setting. Within each column, the rows represent: 1:1 ROC (1st row), 1:1 Top-K Precision (2nd row), 1:10 PRC (3rd row), and 1:10 Top-K Precision (4th row). Observations: In the Standard setting (A–D), GNNs demonstrate robust performance regardless of data balance. Notably, in the challenging Cold-Start (E, F) and Hard-Negative (I, J) scenarios, GNNs maintain acceptable predictive capability under balanced (1:1) conditions. However, a catastrophic performance collapse is observed specifically in the realistic, highly imbalanced (1:10) settings (G, H for Cold-Start; K, L for Hard-Negative). This indicates that while GNNs can handle structural difficulties or class imbalance individually, they fail to generalize when simultaneously confronted with both topological sparsity/bias and realistic negative-sample dominance.

**Table S1. Benchmark comparison on ChemProt (MCQ) and BioASQ (open-ended QA) between open baseline models and our outcome-aligned (SFT and KTO) model.**

| Model name | Chemprot | | | BioASQ | | | | |
|---|---|---|---|---|---|---|---|---|
| | Accuracy | Macro-F1 | Weighted-F1 | GPT score | Rouge-1 | Rouge-2 | Rouge-L | BERTScore-F1 |
| DeepSeek-R1-Distill-Qwen-1.5B | 0.46 | 0.15 | 0.51 | 2.4050 | 0.2513 | 0.0785 | 0.1920 | 0.5501 |
| DeepSeek-R1-Distill-Llama-8B | 0.37 | 0.30 | 0.39 | 3.2220 | 0.2323 | 0.0698 | 0.1765 | 0.5174 |
| DeepSeek-R1-Distill-Qwen-14B | 0.43 | 0.36 | 0.46 | 3.6490 | 0.2723 | 0.0934 | 0.2078 | 0.5445 |
| DeepSeek-R1-Distill-Qwen-32B | 0.44 | 0.34 | 0.47 | 3.7320 | 0.2558 | 0.0869 | 0.1962 | 0.5259 |
| DeepSeek-R1-Distill-Llama-70B | 0.36 | 0.31 | 0.35 | 3.7780 | 0.2769 | 0.0981 | 0.2102 | 0.5555 |
| DeepSeek-R1-671B | 0.48 | 0.39 | 0.50 | 3.9120 | 0.1976 | 0.0547 | 0.1560 | 0.4638 |
| Qwen3-1.7b | 0.24 | 0.22 | 0.23 | 3.4200 | 0.2712 | 0.0889 | 0.2054 | 0.5601 |
| Qwen3-4b | 0.30 | 0.28 | 0.32 | 3.4940 | 0.2657 | 0.0891 | 0.2037 | 0.5544 |
| Qwen3-8b | 0.21 | 0.23 | 0.16 | 3.5100 | 0.2470 | 0.0748 | 0.1884 | 0.5432 |
| Qwen3-14b | 0.65 | 0.46 | 0.68 | 3.6410 | 0.2724 | 0.0914 | 0.2075 | 0.5560 |
| Qwen3-30b-a3b | 0.46 | 0.39 | 0.49 | 3.6360 | 0.2763 | 0.0964 | 0.2114 | 0.5637 |
| Qwen3-32b | 0.27 | 0.27 | 0.23 | 3.6630 | 0.2688 | 0.0877 | 0.2007 | 0.5533 |
| Qwen3-235b-a22b | 0.51 | 0.39 | 0.55 | 3.7300 | 0.2712 | 0.0885 | 0.2029 | 0.5565 |
| Qwen3-8b-kto | 0.24 (+14%) | 0.23 (+0%) | 0.24 (+50%) | 4.0520 (+15%) | 0.2420 | 0.0862 | 0.1739 | 0.6507 (+20%) |

Caption: Benchmark comparison on BioASQ and ChemProt. Our 8B parameter model, aligned with clinical outcomes, achieves state-of-the-art scores on BioASQ factuality metrics, outperforming models orders of magnitude larger.

**Table S2. Ablation across base, SFT checkpoints, and KTO checkpoints on ChemProt and BioASQ.**

| Model name | Chemprot | | | BioASQ | | | | |
|---|---|---|---|---|---|---|---|---|
| | Accuracy | Macro-F1 | Weighted-F1 | GPT score | Rouge-1 | Rouge-2 | Rouge-L | BERTScore-F1 |
| Qwen3-8b | 0.21 | 0.23 | 0.16 | 3.5100 | 0.2470 | 0.0748 | 0.1884 | 0.5432 |
| Qwen3-8b-sft ckpt-3000 | 0.28 (+33%) | 0.27 (+17%) | 0.30 (+88%) | 4.0430 (+15%) | 0.2510 | 0.0893 | 0.1834 | 0.6415 (+18%) |
| Qwen3-8b-sft ckpt-3500 | 0.28 (+33%) | 0.25 (+8%) | 0.31 (+94%) | 3.9980 (+14%) | 0.2463 | 0.0882 | 0.1832 | 0.6330 (+17%) |
| Qwen3-8b-sft ckpt-4000 | 0.29 (+38%) | 0.26 (+13%) | 0.33 (+106%) | 3.9940 (+14%) | 0.2407 | 0.0889 | 0.1781 | 0.6278 (+16%) |
| Qwen3-8b-sft ckpt-4500 | 0.29 (+38%) | 0.26 (+13%) | 0.32 (+100%) | 3.9990 (+14%) | 0.2430 | 0.0878 | 0.1807 | 0.6279 (+16%) |
| Qwen3-8b-sft | 0.29 (+38%) | 0.26 (+13%) | 0.32 (+100%) | 4.0120 (+14%) | 0.2433 | 0.0876 | 0.1797 | 0.6302 (+16%) |
| Qwen3-8b-kto-ckpt-3000 | 0.25 (+19%) | 0.23 (+0%) | 0.25 (+56%) | 4.0500 (+15%) | 0.2410 | 0.0840 | 0.1753 | 0.6470 (+19%) |
| Qwen3-8b-kto-ckpt-3500 | 0.24 (+14%) | 0.23 (+0%) | 0.24 (+50%) | 4.0520 (+15%) | 0.2420 | 0.0862 | 0.1739 | 0.6507 (+20%) |
| Qwen3-8b-kto-ckpt-4000 | 0.24 (+14%) | 0.23 (+0%) | 0.24 (+50%) | 4.0190 (+15%) | 0.2380 | 0.0849 | 0.1716 | 0.6473 (+19%) |
| Qwen3-8b-kto | 0.23 (+9%) | 0.23 (+0%) | 0.23 (+44%) | 4.0250 (+15%) | 0.2392 | 0.0856 | 0.1718 | 0.6500 (+20%) |

Caption: Ablation analysis across training stages. SFT provides the primary boost in mechanistic classification (ChemProt), while KTO further refines generation quality and factuality (BioASQ).

### Supplementary Note 1: Outcome-Aligned KG-RAG Repurposing Framework

```
Input: Raw corpora D0, Knowledge Graph G
Output: Outcome-aligned model parameters θ

// Stage 1: Reasoning-Augmented Data Generation
Initialize D_reason ← ∅
for each (Q, A) in D0 do
    E_Q, E_A ← EntityLinking(Q, A, G)
    Π ← PathRetrieval(G, E_Q, E_A)
    Π_ranked ← SemanticPruning(Π, Q, A)
    C ← GenerateCoT(Q, A, Π_ranked)
    if SelfConsistencyCheck(C, A) is True then
        D_reason ← D_reason ∪ {(Q, C, A)}
    end if
end for

// Stage 2: Supervised Fine-Tuning (SFT)
θ ← InitializeBackbone()
D_SFT ← FilterSuccessful(D_reason)  // Only use approved drugs for SFT
θ_ref ← TrainSFT(θ, D_SFT, Objective=NLL)

// Stage 3: Clinical Alignment (KTO)
// Note: KTO is typically offline. We use pre-generated outputs y with labels o.
D_KTO ← {(Q, y, o)} // y is model generation, o is clinical label (1=Success, 0=Fail)
θ ← θ_ref
for epoch = 1 to E_KTO do
    for batch B ⊂ D_KTO do
        L_batch ← 0
        for (Q, y, o) in B do
            // Compute log-probabilities under current policy and reference
            log_p_θ ← Forward(θ, Q, y)
            log_p_ref ← Forward(θ_ref, Q, y)

            // Likelihood ratio
            log_rho ← log_p_θ - log_p_ref

            // Sigmoid confidence transform
            p ← Sigmoid(κ * log_rho)

            // Binary preference loss (maximize p if o=1, minimize if o=0)
            L_item ← - [ o * log(p) + (1-o) * log(1-p) ]
            L_batch ← L_batch + L_item
        end for
        θ ← Update(θ, ∇ L_batch)
    end for
end for
return θ
```
